# Selection of nanobodies against liponanoparticle-embedded membrane proteins by yeast surface display

**DOI:** 10.1101/2025.05.20.655075

**Authors:** Greg J. Dodge, Hayley L. Knox, Brian Cho, Barbara Imperiali, Karen N. Allen

## Abstract

Single-domain antibodies, known as nanobodies (Nbs), are widely used in structural biology, therapeutics, and as molecular probes in biology and biotechnology. Nbs towards soluble proteins are routinely developed via alpaca immunization or directed evolution in yeast cell-surface display. However, for membrane proteins, the targets are generally detergent-solubilized, and there remains a need for Nb development methods against membrane proteins in a native-like membrane environment. To address this need, we present a protocol for Nb selection via extraction of membrane proteins into amphiphilic polymers such as styrene-maleic acid to produce purified membrane proteins in stable liponanoparticles. Proof of generality is demonstrated by applying the pipeline to four membrane-resident enzymes of differing fold, oligomerization state, and membrane topology (reentrant membrane helix, transmembrane, membrane-associated). Following screening for optimal stabilization into liponanoparticles, Nbs were selected against four target proteins from glycoconjugate biosynthesis pathways. The selected Nbs showed high affinity and selectivity towards their target proteins with K_D_ apparent values ranging from 15 nM to 200 nM, depending on the Nb-protein conjugate. In accordance with their tight binding, various Nb-protein complexes were found to be stable to size-exclusion chromatography purification. The Nbs were also amenable to sortase-mediated ligation, enabling their conversion into molecular probes for the target membrane protein. The ability to select for such high-affinity Nb against membrane proteins in SMALP will facilitate their widespread application in cell biology and biomedical applications.

## Introduction

Antibodies are ubiquitous tools across the life sciences, due to their selectivity and specificity towards discrete protein epitopes and compatibility with functionalization for detection strategies. The epitope recognition region in canonical mammalian antibodies comprises a variable light chain (V_L_) and a variable heavy chain (V_H_). Unfortunately, the oligomeric nature and obligate glycosylation of such antibodies complicates production and prevents large-scale economic expression in bacteria. In contrast to the typical mammalian antibody scaffold, that of camelids such as llama and alpaca has evolved to include an epitope recognition surface that comprises solely a heavy chain (V_HH_).^1^ These single V_HH_ domains, referred to as nanobodies (Nbs), can be expressed and purified in multi-milligram quantities from *Escherichia coli*.^2^ As nanobody (Nb) domains retain the binding and specificity characteristics of antibodies, but are smaller (∼15 kDa vs. ∼150 kDa) and can be produced more easily, they have been employed widely as research tools in cell biology, as well as medical applications.^3^ For example, Nb-based therapeutics have become increasingly deployed in recent years. A range of novel modalities have been adapted to include V_HH_ moieties for delivery and targeting, including Nb-CAR-T, Nb-liposomes, and Nb-drug conjugates.^4^ Two Nb-based therapies have successfully advanced through clinical trials, gaining approval by the FDA for treatment of acquired thrombotic thrombocytopenic purpura^5^ and relapsed or refractory multiple myeloma,^6^ with > 12 additional Nb therapeutics in various stages of clinical trials.^4^

In addition to their promise as therapeutics and molecular probes, nanobodies have revolutionized membrane protein structural biology. Landmark studies on G protein-coupled receptors (GPCRs), specifically the β_2_-adrenergic receptor, have demonstrated the utility of nanobodies as both crystallization chaperones and reagents to trap receptors in specific conformations.^7–8^ More recently, nanobodies have played a central role in strategies for sample stabilization and the addition of fiducial mass markers for Cryo-EM.^9–11^ By utilizing Nb-mediated complex formation, Cryo-EM reconstructions of small proteins such as the 26 kDa KDEL receptor are possible,^9^ a result which would otherwise be beyond the scope of current methodologies. These advances clearly demonstrate the utility of small, high-affinity binders for the greater structural biology community.

Although nanobodies hold great potential as both molecular probes and tools for structural biology, methods for their rapid generation have lagged. Previously, most Nb selection campaigns relied on active immunization of primary organisms, such as llama, with a protein of interest followed by cloning of the antibody variable region from isolated sera to generate the library.^2^ This workflow is particularly challenging for membrane proteins, which may suffer from very low yields using the typical approach of recombinant expression, solubilization in detergent and purification, making it challenging to produce the required quantities of protein for inoculation. To address this limitation, alternative approaches to active immunization have been developed, including peptide or protein display on phage, yeast, and ribosomes.^12–14^

Concomitantly, strategies for production and purification of integral membrane proteins have been advancing. One area of active development is the use of amphiphilic polymers such as styrene maleic acid (SMA) to extract proteins and surrounding lipid directly from cell membranes in liponanoparticles (SMALPs).^15–16^ The SMALP-based methodologies are advantageous relative to classical detergent-extraction techniques, as target proteins remain in a lipid bilayer throughout the purification process, preserving both stability and native lipid interactions. SMALP-solubilized proteins are generally more stable when compared to detergent-solubilized targets and in some cases, SMALP solubilization significantly increases yields of target membrane proteins ^16–17^ We have recently described an optimized expression and purification scheme to generate bacterial membrane proteins in SMALPs for structural biology applications.^18^ This protocol was applied to a single example within the monotopic phosphoglycosyl transferases (PGTs) superfamily from prokaryotes; however, the generalizability of the method remained unclear. Despite the utility of SMALPs for retention of membrane proteins in a native-like environment, the general utilization of SMALPs for Nb selection remains unexplored.

Here, we establish the generality of the method by applying it to prokaryotic proteins of diverse membrane topologies. Leveraging the dual-Strep tag and as described in the original example,^18^ a yeast-surface display protocol to select Nbs against membrane proteins in SMALP is presented. The successful Nb-SMALP candidates were fully characterized by applying complementary biophysical methods. In addition, the application of sortase-mediated ligation (SML)^19^ is demonstrated for site-specific Nb functionalization, providing a facile route for converting selected Nbs into molecular probes.

## Results & Discussion

The utility of Nbs is undeniable as shown by their widespread use as tools, probes and therapeutics.^3^ However, their application to the important category of membrane proteins is hampered due to limited compatibility between Nb selection strategies and membrane protein solubilization strategies. To overcome this obstacle, a protocol for membrane protein extraction and purification in polymer-stabilized liponanoparticles was applied, followed by Nb selection by yeast-surface display. The generality of this protocol was demonstrated by applying it to membrane proteins of differing fold and membrane topology.

### Purification of bacterial membrane proteins in SMALP

To establish a general protocol based on the method used for purification of the Lg-PGT WbaP from *Salmonella enterica* O-antigen biosynthesis in liponanoparticles^20^ a subset of membrane proteins from glycoconjugate biosynthesis pathways with varying membrane topology, size, oligomerization, and fold were selected (**Figure 1A**, **Table 1**). Following the same screening protocol applied to WbaP, purification conditions for all targets were identified utilizing a small collection of SMA co-polymers comprised of SMALP 140, SMALP 200, SMALP 300, diisobutylene maleic acid (DIBMA).^21^ For selected cases (Table 1) AASTY 6-45 and AASTY 6-50 were added to the set.^22^ Most target proteins preferred SMALP 200 (previously called SMA30) polymer, except for PglI from *Campylobacter concisus*, which preferred SMALP300 (previously SMA40). Owing to the resulting high purity and yield of these targets (**Figure S1)**, this set of proteins, along with the previously reported WbaP, were used as candidates for Nb selection.

**Figure 1.**
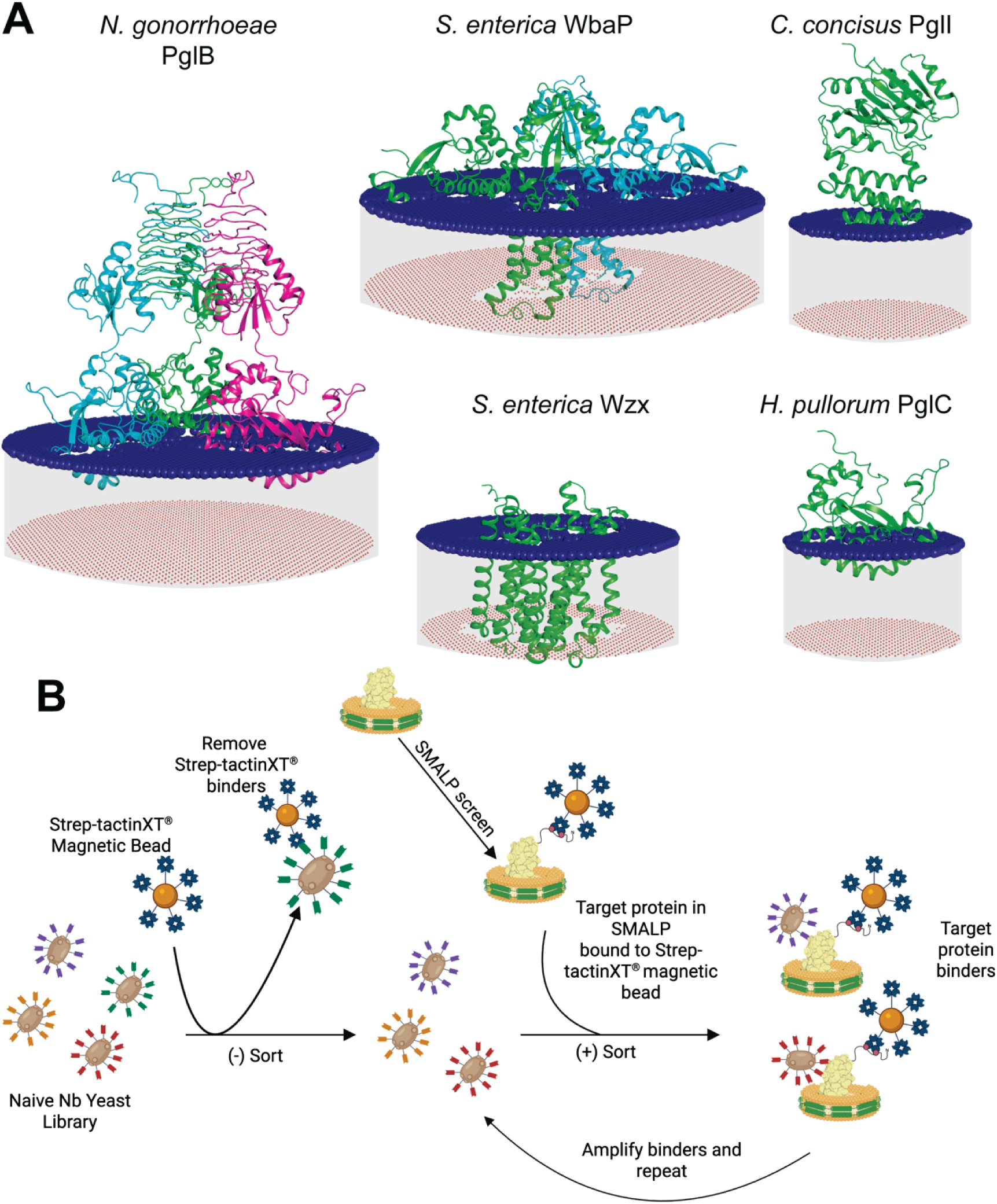
Target proteins and nanobody selection strategy. (**A**) Selected prokaryotic membrane proteins, generated in AlphaFold (apart from WbaP wherein PDB ID: 8TB3 was used) and modeled into membranes (represented by blue and red spheres) using the PPM3 server. (**B**) Yeast-surface display strategy for selection of binders towards target proteins.

**Table 1.** Selected targets for Nb screen.

| **Protein** | **Organism** | **UniProt ID** | **Protein superfamily/family** | **Protein Fold** |
| --- | --- | --- | --- | --- |
| WbaP | *Salmonella enterica* (LT2) | P26406 | Monotopic PGT/Large PGT | PGT |
| PglC | *Helicobacter pullorum* | E1B268 | Monotopic PGT/Small PGT | PGT |
| Wzx (RfbX) | *Salmonella enterica* (LT2) | P26400 | Polysaccharide biosynthesis and transport | Transmembrane helices |
| PglB | *Neisseria gonorrhoeae* (strain ATCC 700825 / FA 1090) | Q5FAE1 | Monotopic PGT/Bifunctional | Acetyltransferase/trans-hexapeptide repeat/PGT |
| PglI | *Campylobacter concisus* strain 13826 | A7ZEU0 | Nucleotide-diphospho-sugar transferases/Glycosyltransferase-2 | Rossmannoid |

### Selection of nanobodies against purified bacterial membrane proteins by yeast display

After assessing the various methods available for selection of Nbs, the yeast surface display (YSD) method developed by McMahon and coworkers was adopted.^12^ The rationale behind this choice was: (1) commercial availability of the library; (2) low cost and ease of culture for yeast; and (3) amenability of the selection protocol to modification. In the original protocol the selection relied on non-specific labeling of lysine residues on target proteins with fluorophores bearing an N-hydroxysuccinimide (NHS) ester, followed by sorts with anti-fluorophore antibody magnetic beads or fluorescence-activated cell sorting (FACS). Although this protocol was quite successful for the original G protein-coupled receptor (GPCR) targets, there are several technical downsides. Notably, the use of NHS-fluorophores limits control over labeling of target proteins and the potential over-functionalization may enrich fluorophore binders. We hypothesized that the dual-Strep tag used for purification of target proteins could serve an additional role as a handle for selection in YSD, as both streptactin-functionalized magnetic beads and fluorophore-conjugated streptactin are commercially available. Therefore, a modified workflow was developed for YSD selection, which does not require functionalization of target proteins (**Figure 1B**).

The initial target chosen to benchmark and optimize YSD Nb selection against proteins in SMALP was WbaP. This protein is well characterized, stable, and can be purified in milligram-scale from recombinant expression in *E. coli* (**Figure S1**).^20^ After expanding and expressing the yeast library, the initial selection relied on an iterative sequence involving two negative sorts followed by a single positive sort. The first negative sort was against empty MagStrep® Strep-Tactin® XT beads (Strep-Tactin XT beads) and served to remove bead and streptactin binders. The second negative sort was against Strep-Tactin XT beads loaded with WecP, a WbaP homolog from *Aeromonas hydrophila,* purified in an identical manner to WbaP in SMALP 200 liponanoparticles (**Figure S2**). This step served to remove binders to the dual-Strep tag, or any of the components of the liponanoparticle.

Enrichment of the yeast population for WbaP-binding was monitored via analytical flow cytometry (**Figure 2**). After three rounds of magnetic assisted cell sorting (MACS), 8.0 % of cells resulted from the final round of binder selection, an increase from 0.7 % in the naïve library (**Figure 2**). We next assessed the specificity of our selected populations for WbaP over the homolog WecP. Although there was a clear increase in binding over the course of the MACS selections for WbaP, no such enrichment was observed for WecP (**Figure S2**). This result demonstrates that despite the compositional complexity of the protein-laden SMALP, binders were selected in a protein-specific manner. To further challenge the binding capability of selected yeast, the concentration of WbaP was reduced from 500 nM to 250 nM and cells were sorted using fluorescence-activated cell sorting (FASCS) to generate population 0.4 (**Figure S3**). To select for binders specific to the N-terminal membrane-associated domains of WbaP, a truncated version of the protein was purified in SMALP and used for the final sort. After one round of FACS, population 0.4 comprised 18 % positive WbaP binders, even at the lower 250 nM WbaP concentration (**Figure 2A**).

**Figure 2.**
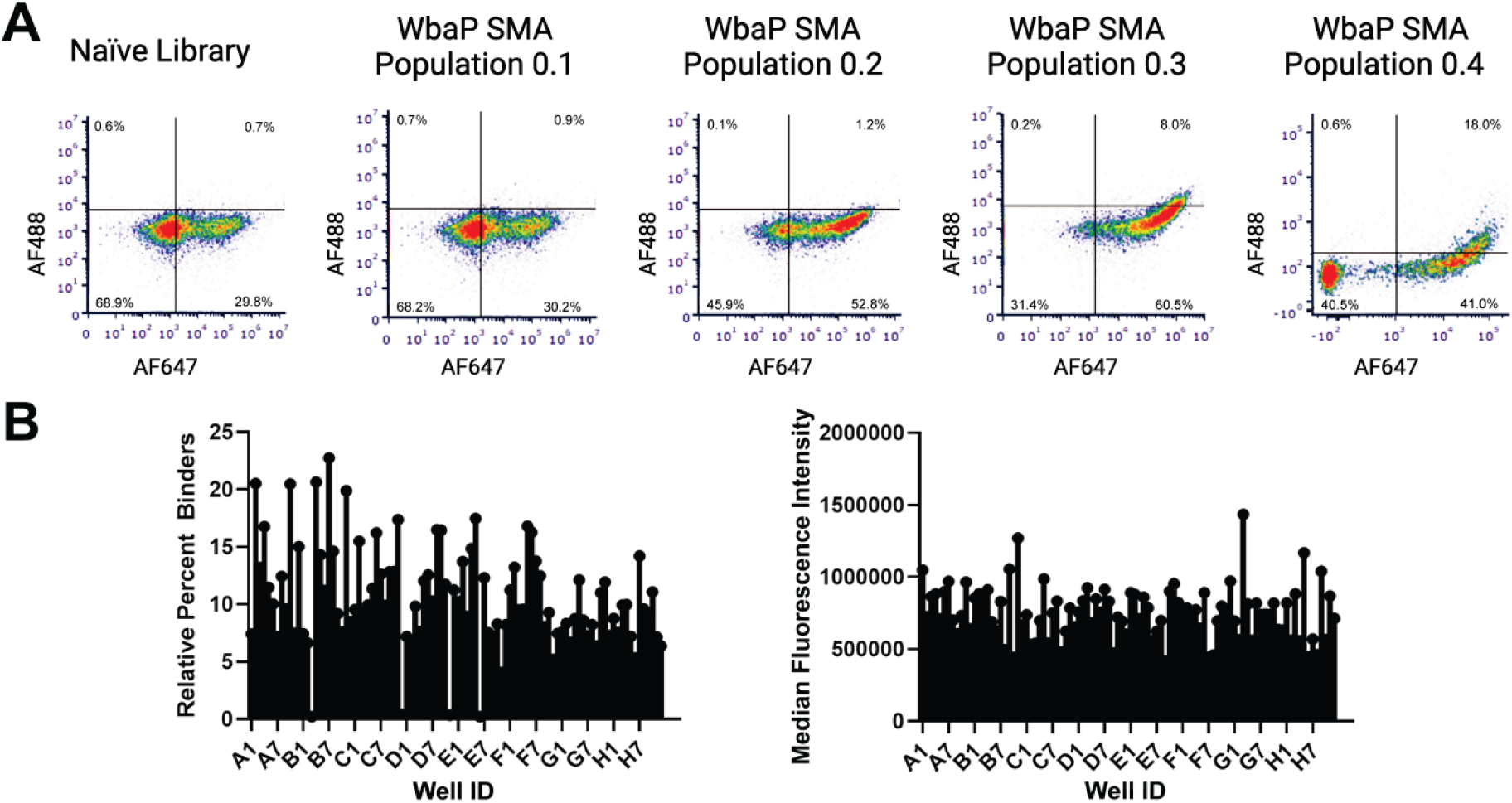
Enrichment of binders against *S. enterica* WbaP in SMALP (**A**) Analytical flow cytometry analysis of successively selected populations. After each round of sorting, the percentage of cells in the upper right quadrant (WbaP binders) increased. Populations up to 0.3 stained using 500 nM WbaP, population 0.4 stained using 250 nM WbaP. (**B**) Analysis of selected individual clones in a 96-well plate from WbaP SMA population 0.4 based on (left) relative percent binders and (right) median fluorescence intensity. 100 nM WbaP used for staining.

Prior to single-clone analysis, population 0.4 was challenged with a serial dilution of WbaP from 1 μM to 1 nM to identify the optimal concentration for binding analysis by analytical flow cytometry. The optimal mean fluorescence intensity (MFI) was observed at 100 nM, with sensitivity down to 1 nM (**Figure S4**). This result is consistent with a ‘hook effect’ in which over-saturation of a component in a binding experiment results in reduction of signal.^23^ In a 96-well plate format individual colonies were isolated from population 0.4 and used to identify the top binding Nb clones against 100 nM WbaP using analytical flow cytometry (**Figure 2B**).

With the success of the binder selection for WbaP, binders were selected against the membrane proteins from the expanded SMALP purification panel (Table 1). After three rounds of MACS and a single round of FACs, binders of *S. enterica* Wzx (formerly referred to as RfbX), *N. gonorrhoeae* PglB, and *C. concisus* PglI were selected. However, we were unable to enrich for binders against *H. pullorum* PglC purified in diisobutylene maleic acid liponanoparticles (DIBMALP). This likely reflects poor quality of PglC purified in DIBMALP and is discussed below. Ninety-six colonies isolated from the final populations of each target were used to identify top binding single Nb clones (**Figure S5**).

### Biophysical characterization of Nbs

Sequence analysis demonstrated that the top binder selected for each target was unique, with a maximum sequence identity between Nbs of ∼86% (**Figure S6**). Top binding Nb sequences identified for each target were cloned into a pET22b vector for periplasmic expression in *E. coli*.^24^ Each Nb was isolated to high yield and purity after recombinant expression (**Figure S7**). To investigate the stability of the purified nanobodies we utilized nanoscale differential scanning fluorimetry (nanoDSF). For each of the purified Nbs, thermal transitions were observed to be > 70 °C (**Figure S8**), demonstrating that the synthetic Nbs from the yeast library are stable. We then assessed direct binding between purified Nbs and their targets using biolayer interferometry (BLI). For this assay, Nbs were immobilized onto NiNTA functionalized BLI tips via the C-terminal His_6_ tag used for purification. Association and dissociation curves were observed for all target membrane proteins, with K_D_ (apparent) values ranging from ∼200 nM to ∼15 nM depending on the Nb – protein pair (**Figure 3**).

**Figure 3.**
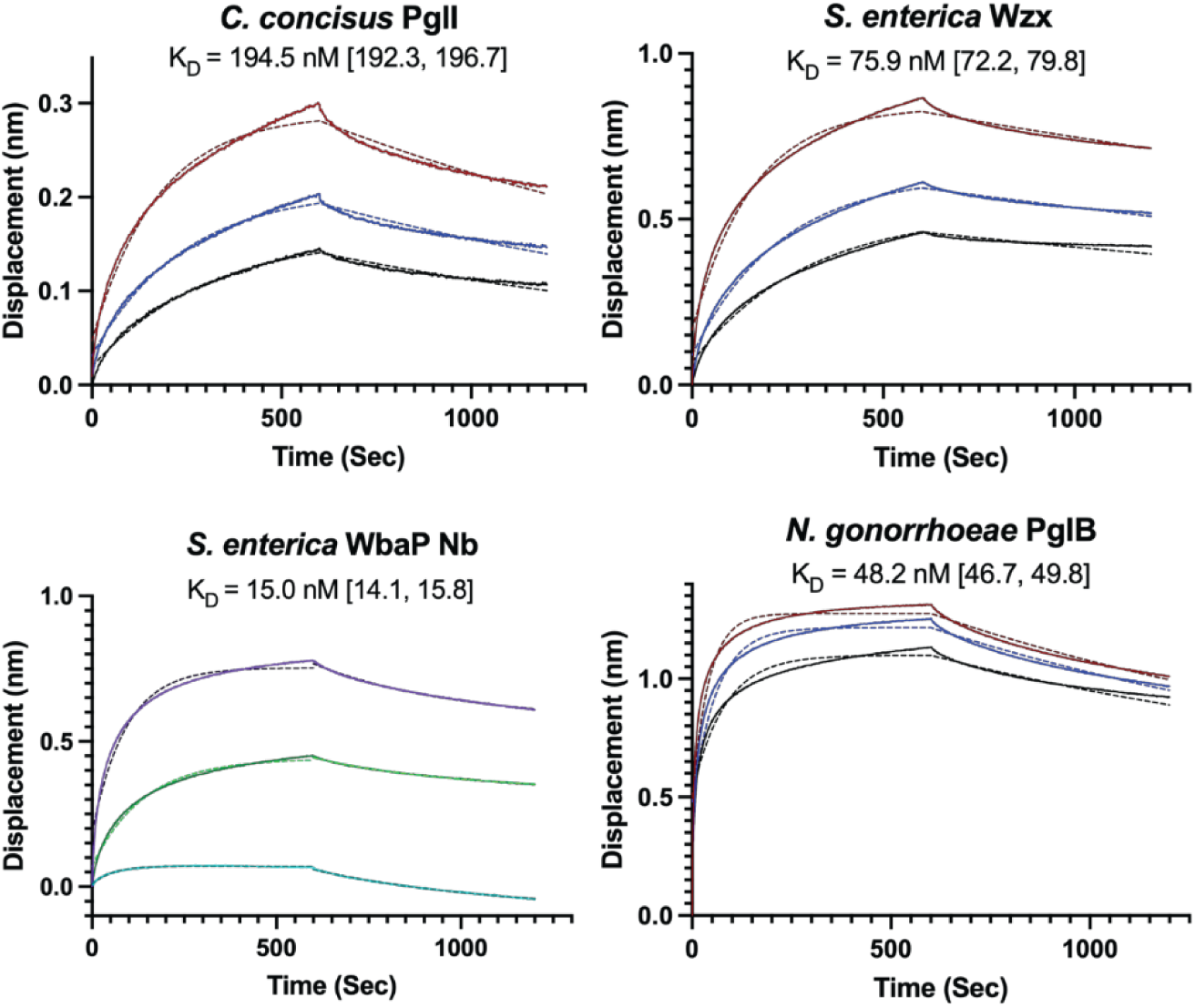
Binding data and coefficients for Nbs and their cognate protein with 95% confidence intervals in brackets. Protein concentrations are 1500 nM (red), 1000 nM (blue), and 750 nM (black) for PglI, Wzx, and PglB Nbs. Protein concentrations are 500 nM (purple), 250 nM (green), and 125 nM (teal) for WbaP Nbs. All curves were done in duplicate. Solid lines indicate the collected data and the fit is in dashed lines. For the WbaP Nbs, the 1-phase association and dissociation curves were fit separately. All K_D_ values are apparent dissociation constants.

### Functionalization of nanobodies using sortase mediated ligation (SML)

Sortase mediated ligation (SML) is a common technique for conversion of protein binders into molecular probes (**Figure S9**).^19, 25^ This technique relies on the sortase A recognition of an LPETG sequence motif. Upon motif recognition, sortase A cleaves the peptide backbone between the T and G amino acids, forming a covalent thioester intermediate with the protein bearing the motif. Sortase A will then couple the N-terminus of any peptide bearing an N-terminal GGG motif to the C-terminus of the protein to be sortagged.

Functionalized polyglycine peptides are accepted as sortase substrates, allowing for site-specific enzymatic transfer of biotin or fluorophores onto a protein of interest. SML has recently been used to make molecular probes from human lectins and to generate biotinylated Nbs.^19, 26^ Modified pET22b vectors were constructed to insert an LPETG motif between the Nbs and C-terminal His_6_ tags. Purified sortagged Nbs were then functionalized using a GGGYK-SulfoCy5 peptide as previously reported.^26^ Confirmation of Nb labeling was accomplishing via SDS-PAGE, with gels imaged in the Cy5 channel prior to staining with Coomassie dye, showing the Cy5 peptide was successfully transferred (**Figure S10**).

A variant of native gel electrophoresis, SMA-PAGE,^27^ was used to monitor formation of complexes between Cy5-labeled Nbs and their target proteins. In this method, the charge from the SMA polymer allows protein-loaded SMALPs to migrate along the voltage gradient without destruction of the SMALPs or denaturation of protein. Clear migration of both WbaP and Wzx was observed (**Figure S11**), but not for PglB or PglI (data not shown). We hypothesize that the differences in native gel migration may be due to differences in the membrane embeddedness of the targets. The PPM server^28^ was used to predict the ΔG transfer free energies of the targets into membrane. Both WbaP (ΔG =-123.7 kcal/mol) and Wzx (ΔG = - 98.9 kcal/mol) are predicted to have significant ΔG transfer free energies into the membrane, whereas PglI (ΔG =-16.9 kcal/mol) and PglB (ΔG =-46.8) have less of their mass embedded in membrane. As the isoelectric point (pI) of proteins strongly influences migration in native PAGE and all these proteins have pI ≥ 8.0, the more hydrophilic nature of PglB and PglI may prevent migration, regardless of the net charge contributed from the SMA polymer. Likewise, we see minimal signal for Cy5-labeled Nbs alone when monitored by fluorescence or by Coomassie staining. However, samples of WbaP and Cy5-Nb or Wzx and Cy5-Nb show fluorescent bands on the gel and Coomassie staining in these regions, indicating that additional fluorescent Nb is co-migrating with membrane proteins in SMALP.

### Stability of Nb complex assessed by size-exclusion chromatography

Size-exclusion chromatography (SEC) has been used to verify complexation between Nbs and their target proteins for crystallization and cryo-EM sample preparation. Importantly, as the injected material is highly diluted over the column resin bed, certain complexes may dissociate. For testing the complex stability, we chose both WbaP and PglI with their cognate Nbs. For both targets, the complexes remained intact and a significant shift in elution volume was observed for the complex relative to a control injection lacking Nb (**Figure 4**). Following this, we pursued a maltose-binding protein fusion to the Nb, referred to as a promacrobody (PMB),^29^ which results in a larger construct that is of a mass more suited for CryoEM. WbaP and PglB were chosen for this analysis due to the differences in their membrane topologies. The complexes remained intact as assessed by SDS-PAGE. For both WbaP and PglB, shifts are seen in the elution volumes relative to controls lacking PMB. Both Nb and PMB fusions were demonstrated to show tight binding to their cognate proteins which results in a pure complex that can be used for further structural characterization. Importantly, two of the enzymes, PglI and PglB, were isolated in a different polymer than the one used for Nb generation and Nb and PMB complexation was still observed, confirming that the Nbs are specific for the protein of interest, not the polymer.

**Figure 4.**
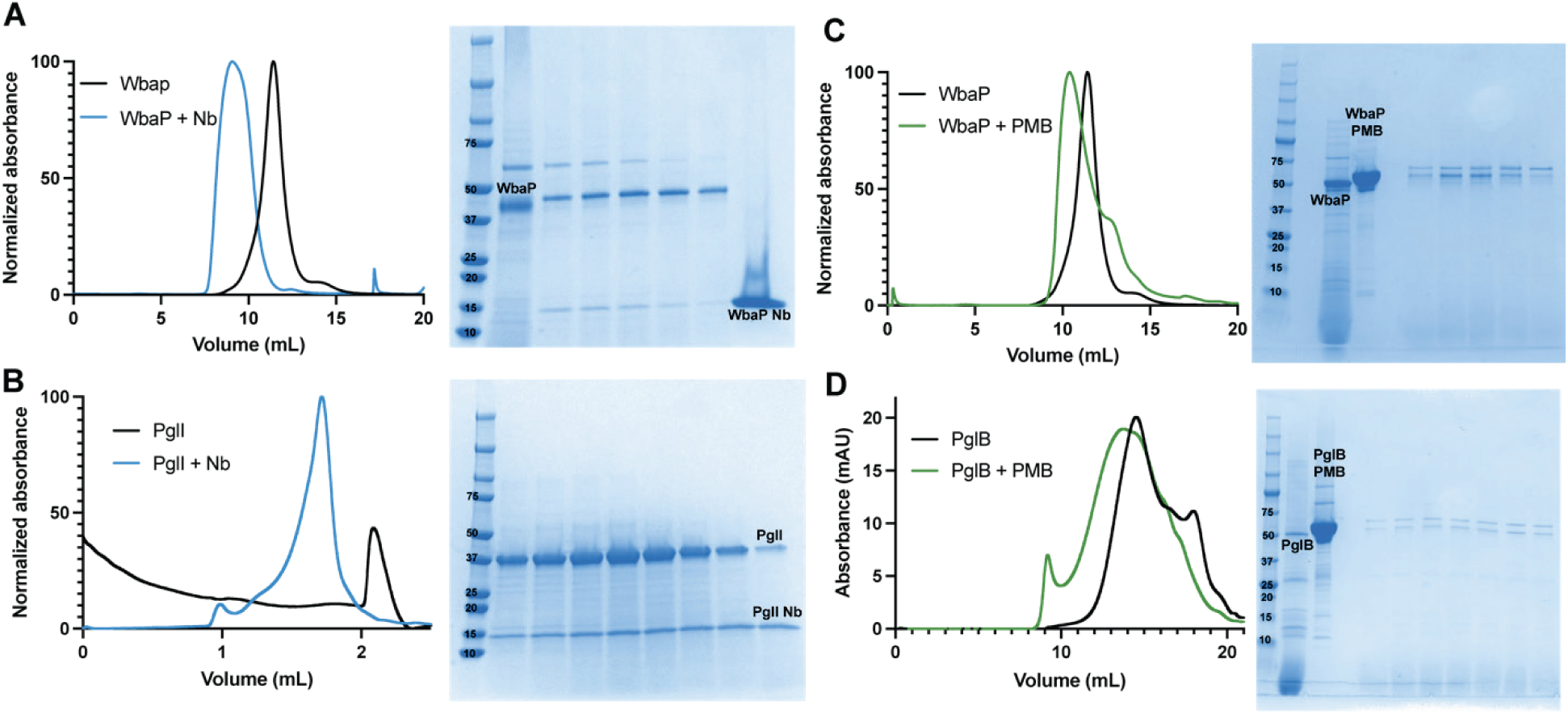
SEC data and SDS-Page of selected enzymes with their cognate Nbs. (**A**) WbaP with Nb, sized on an Enrich 650 10/300 column. (**B**) PglI with Nb, sized on a Superose 6 Increase 3.2/300 column. (**C**) WbaP with PMB, sized on an Enrich 650 10/300 column. (**D**) PglB with PMB, sized on a Superose 6 10/300 GL column.

## Conclusions

Herein, we describe a modified yeast-surface display protocol compatible with Nb selection against membrane proteins embedded in native-like liponanoparticles. We remove the limiting requirement of fluorescently labeling target membrane proteins prior to YSD selection used in established protocols through utilization of the same dual-Strep tag for both purification and immobilization on magnetic beads during the selections. This methodology is compatible with SMALP-embedded targets, which significantly expands the utility for the study of membrane proteins. SMALPs often confer enhanced stability for membrane proteins and additionally can sample more native-like conformations due to the similarity to the lipid bilayer. Additionally, some membrane proteins are recalcitrant to solubilization in detergent and this property limits biochemical and biophysical applications. Notably, our results indicate that this methodology is applicable to a wide variety of membrane protein topologies. Although the targets are currently limited to prokaryotic membrane proteins, we anticipate the methodology described here will also be applicable to eukaryotic membrane proteins.

The selected Nbs are stable, readily produced recombinantly in *E. coli,* and bind specifically to their targets as validated using a variety of complementary methods. Experimental K_D_ (apparent) values support that selected Nbs have high affinities for their cognate antigens, even in the absence of affinity maturation. We also observe the Nb-antigen complex either by SMA-PAGE or SEC, showing the stability of the complex. In the case of SEC, the complex is ready for further structural studies, such as X-ray crystallography or CryoEM. Additionally, through the coopting of the sortase A reaction and recognition motif, selected Nbs can be efficiently converted into fluorescent molecular probes. This opens the possibility of in-house production of reagents for detection of specific membrane protein antigens in a variety of experiments, such as western blotting and fluorescence microscopy. Furthermore, we anticipate the Nbs selected here will play central roles in structural characterization of the PglB, PglI, and Wzx targets, which have no reported experimental structures. Specifically, we envision using Nbs to conformationally stabilize targets and act as tools for addition of fiducial mass in CryoEM experiments.

More broadly, the ability to rapidly select high-affinity binders against target membrane proteins in SMALP has implications for both vaccine design and delivery of therapeutic molecules. The MACS protocol described herein requires picomoles of material, which allows for the study of targets with low purification yields. Finally, we have shown that the selected Nbs are compatible with fluorophore functionalization for probe generation. This methodology can be coopted to fuse therapeutic small molecules, proteins, or larger cargo to Nbs for targeted delivery to specific membrane proteins, cell types, or tissues. The methodology herein extends the functionality of Nbs as probes of membrane proteins, an expanding methodology in varied contexts.

## Materials and Methods

### Membrane protein methods

#### Cloning, Expression and purification

Gene fragments encoding target proteins were codon optimized for expression in *E. coli* and inserted into a SUMO-strep-TEV vector using Gibson assembly as described previously (**Table 1**).^18^ A truncated construct encoding residues 1-272 of WbaP was ordered as a synthetic gene block, and inserted into a SUMO-strep-TEV vector using Gibson assembly.

Expression and purification of proteins followed a previously established protocol.^18^ Briefly, C43 *E. coli* cells harboring plasmid pAM174^30^ were transformed with plasmid encoding a target protein. Transformed cells were used to inoculate 0.5 L terrific broth autoinduction^31^ cultures supplemented with 150 μg/mL kanamycin and 25 μg/mL chloramphenicol. Cultures were incubated at 37° C until OD_600_ measured ∼1.5, at which point the temperature was adjusted to 18° C and 1 g solid L-arabinose was added. After ∼20 hours of expression, cells were isolated via centrifugation, transferred to a 1-gallon Ziplock® bag, and frozen at - 80° C.

For purification, frozen cells were resuspended in buffer A (50 mM HEPES pH 8.0, 300 mM NaCl) to a final volume of 4 mL per g cell pellet. Resuspended cells were supplemented with 2 mM MgCl_2_, 0.06 mg/mL lysozyme (RPI) and 0.5 mg/mL DNase I (Millipore Sigma) and incubated on ice for 30 minutes. Cells were lysed using an M110P microfluidizer (Microfluidics) at 18,000 PSI. Cell membranes (CEF) were isolated by differential centrifugation using a Ti-45 rotor at 9000g for 45 min followed by centrifugation of the supernatant at 160,000g for 65 min. Isolated cell membranes were resuspended to a final concentration of 50 mg/mL in buffer A based on UV absorbance at 280 nm, frozen in liquid N_2_, and stored at-80° C.

WbaP, Wzx and PglC were screened against a small library of styrene maleic acid copolymers to identify optimal liponanoparticle (SMALP) solubilization parameters.^18^ Polymers screened were SMALP 140, SMALP 200, SMALP 300 (Orbisope), as well as diisobutylene maleic acid (DIBMA) (BASF). Several aliquots of 4.5 mL frozen CEF were thawed, mixed 1:1 with a 2% stock solution of each of the polymers in Buffer A, and rotated at room temperature for one hour. Soluble SMALPs were isolated via centrifugation (Ti-45 rotor, 160,000g for 65 min). The supernatant was flowed over 0.5 mL Strep-Tactin®XT 4Flow® resin (Streptactin 4flow, IBA Biosciences) pre-equilibrated with buffer A. The resulting flow through was applied to the column a second time, and the column was washed with 5 column volumes (CVs) buffer A. Immobilized SMALPs were eluted using 3 CV buffer A supplemented with 50 mM biotin (Buffer B). Fractions were analyzed by SDS-PAGE to identify optimal SMA/DIBMA polymer. After the original screen, alternative polymers were added to the screen to yield high quality PglI and PglB, namely AASTY 6-45 and AASTY 6-50 (Cube Biotech, Germany) respectively, that were used for the assessment of complexation by SEC. Purification methods for these polymers were identical to those for the original screen.

Once optimal solubilization conditions were identified, large-scale purifications of each target using membranes isolated from 0.5 L of culture proceeded as previously described.^20^ Isolated cell membranes were mixed 1:1 with a 2% stock solution of the optimal SMA/DIBMA polymer, and rotated for one hour at room temperature. Soluble liponanoparticles were isolated by centrifugation (160,000g, 65 min), and the supernatant was flowed twice over 1 mL Streptactin 4flow pre-equilibrated with Buffer A. Columns were washed with 5 CVs Buffer A and eluted using 3 CVs Buffer B. Protein containing fractions were pooled, and biotin was removed using 3x tandem HiTrap desalting columns (Cytiva) equilibrated with 25 mM HEPES pH 8.0, 150 mM NaCl (Buffer C). Desalted SMALPs were concentrated to 500 μL using Vivaspin 30 kDa concentrators (Sartorius), and injected onto a 24 mL SEC-650 Enrich column (Biorad) pre-equilibrated with Buffer C. Peak fractions were pooled, concentrated to 1 mg/mL, flash frozen in liquid N_2_, and stored at - 80° C.

### Nanobody (Nb) Selection

For this work, a modified version of the selection protocol utilized by McMahon *et al*^12^ was developed.

#### Naïve library expansion

A commercially available yeast-display Nb library (NbLib, Kerafast) was utilized for all selections.^12^ A vial of 2×10^10^ cells was thawed at 30° C, and used to inoculate 1 L of Yglc4.5-Trp + glucose media. Cells were incubated for 18 hrs at 30° C while shaking at 230 RPM. Resulting cells were used to seed 3 L of Yglc4.5 - Trp + glucose and incubated for 48 hours at 30° C while shaking at 230 RPM. After 48 hrs, 2.8×10^8^ cells per mL was achieved, verified by measuring OD_600_ wherein an OD_600_ of 1 is equal to 1×10^7^ cells. Cells (1 L) were pelleted via centrifugation and resuspended in 28 mL YglC4.5-Trp + 10% DMSO and dispensed in 2 mL aliquots into cryovials to give a final concentration of 2×10^10^ cells per vial.

#### Naïve library expression and analysis

Expression media (1 L of Yglc4.5-Trp + galactose) was inoculated with 2×10^10^ cells and incubated for 72 hours at 25 °C while shaking at 220 RPM. After expression, OD_600_ reached 15.96, corresponding to 1.57×10^8^cells per mL culture.

Analytical flow cytometry was used to assess the level of Nb expression of the induced naïve library. A sample of 1×10^7^ cells from the induced culture were washed 2X in 1 mL of cold phosphate-buffered saline plus 0.1% bovine serum albumin (PBSA), and brought to a final volume of 500 μL, corresponding to 2×10^7^ cells per mL. Following this, 20 μL cells were dispensed into a 96-well V-bottom plate (Corning) and diluted with 80 μL cold PBSA. Immunostaining of cells was accomplished using a primary chicken α-hemagglutinin (HA) antibody (Exalpha) with a corresponding secondary goat α-chicken antibody conjugated with an AlexaFluor™ 647 dye (Invitrogen). Strep-tactin XT binding was assessed using Strep-tactin®XT conjugated to AlexaFluor™ 488 dye (IBA Biosciences). Flow cytometry data were collected on an iQue Screener instrument (Sartorius).

#### Magnetic-assisted cell sorting (MACS)

A 50 μL aliquot of MagStrep® Strep-Tactin® XT beads (Strep-tactin XT beads, IBA Lifesciences) was washed 3x in 1 mL cold PBSA using a magnetic rack (Sergi Lab Supplies) to isolate beads. After the final wash, beads were resuspended, and aliquoted into separate 250 μL aliquots. Each target protein was subjected to the following sorts in order: a negative sort against empty Strep-tactin XT beads, a negative sort against strep-tactin XT beads bound to a control Strep-tagged protein in SMALP, and a positive sort against the desired target protein in SMALP bound to Strep-tactin XT beads. For the control and target protein sorts, beads were loaded with 1 pmol protein for 1 hr at 4 °C, and washed 3x with cold PBSA. The sorts consisted of incubation of magnetic beads with 1 mL of Nb-expressing yeast at ∼1×10^8^ cells per mL for 1 hour at 4°C. After each negative sort, unbound yeast cells were transferred to the subsequent tube and beads were discarded. After the final positive sort, unbound cells were removed, and beads were washed 3x with 1 mL cold PBSA. After the final wash, PBSA was removed, and 1 mL YglC 4.5-Trp + glucose growth media was added. Beads were resuspended in media and serially diluted 10-fold 3x. Following this, 50 µL of each dilution was plated on solid Yglc4.5-Trp agar plates and incubated at 30 °C until colonies were visible, typically after ∼48 hours. Colonies were counted and used to quantify diversity after sorts. The remaining 990 µL undiluted magnetic bead containing media was used to inoculate a 5 mL liquid YglC 4.5-Trp + glucose cultures. Cultures were incubated at 30 °C and 220 RPM overnight. Enough cells to ensure 10x diversity measured on plates was used to inoculate 5 mL-Trp + galactose expression media. Each target was subject to two separate rounds of MACS consisting of the steps described above.

#### Analytical flow cytometry

All sorted populations were assessed by analytical flow cytometry on an iQue Screener instrument (Sartorius). Induced yeast cells were assessed for binding by incubation with a cognate membrane protein in SMALP at various concentrations in the presence of a chicken anti-HA antibody in PBSA. After washing, cells were stained with Alexa Fluor 488-functionalized Strep-tactin®XT (IBA Lifesciences) and goat anti-chicken antibody functionalized with Alexa Fluor 647 (AF647). All data was analyzed using FCS Express (De Novo Software, Glendale CA). Binding curves were computed using GraphPad Prism (GraphPad Software, San Diego, CA). All data were collected and analyzed in triplicate biological replicates.

#### Fluorescence assisted cell sorting (FACS)

After two rounds of MACs, a final sort was accomplished using a FACS Aria II instrument (BD Biosciences, San Jose, CA). Induced yeast cells were washed with cold PBSA, incubated with cognate membrane protein in SMALP and a chicken anti-HA antibody. Cells were washed with cold PBSA, then stained with Alexa Fluor 488-functionalized Strep-tactin® XT (iba biosciences) and AF647 functionalized goat anti-chicken antibody.

#### Clonal yeast selection

Final yeast populations were plated on Yglc-Trp + Glu agar plates and incubated 48 hours at 30°C. 96 individual colonies were picked and transferred to a 96-well block pre-filled with 1 mL Yglc4.5-Trp + glucose liquid media. The block was sealed with an air-permeable filter (Diversified Biotech) and incubated for 48 hours at room temperature shaking at 200 rpm. Clonal yeast was then induced by seeding 25 µL of each individual culture into a 96 well block pre-filled with 1 mL Yglc4.5-Trp + galactose media. Plates were sealed with an air-permeable filter and incubated at room temperature for 48 hours with shaking at 200 rpm. Clonal yeast binding was assessed using analytic flow cytometry. The top binding clones were used to inoculate 5 mL YglC4.5-Trp + glucose media, grown overnight at 30 °C 220 RPM, and plasmid DNA was isolated using the Zymoprep™ Yeast Plasmid Miniprep II kit (Zymo Research). The resulting plasmids were sequenced via Sanger sequencing.

### Nanobody Purification

Single Nb coding regions were amplified from pYDS yeast vectors via PCR (**Table S1**), which add complementary overhangs compatible with the *E. coli* expression vector pET22b (Novagen). Sequences were confirmed using Sanger Sequencing.

pET22b plasmids containing single Nb clones were used to transform BL21 (AI) *E. coli* for expression. A 50 μL sample of freshly transformed cells was used to inoculate 3 mL LB cultures supplemented with 100 μg/mL carbenicillin. Cultures were incubated overnight at 37 °C and 225 RPM, then used to inoculate 0.5 L Terrific Broth^32^ supplemented with 100 μg/mL carbenicillin. Cells were allowed to grow to an OD ∼1 at 37 °C and 225 RPM, after which the temperature was reduced to 18 °C, and cultures were supplemented with 1 g powdered L-arabinose and isopropyl β-D-1-thiogalactopyranoside (IPTG) to a final concentration of 0.2 mM. Cells were harvested via centrifugation, transferred to 1 gallon Ziploc® bags, spread to a thin layer, and frozen and-80 °C.

Nanobodies were purified as described previously.^12^ Purified nanobodies were concentrated to 2 mg/mL, and flash frozen in liquid N_2_, and stored at-80 °C.

#### Generation of Nanobodies for Sortase Mediated Ligation (SML)

A sortase recognition motif^19^ and a short glycine linker (SGGSGSGGLPETG) were added to all pET22b Nb sequences using the Q5® Site-directed Mutagenesis Kit following the manufacturer’s recommended protocol. The overlapping primers universally compatible with the nanobodies from the Nb Library (**Table S1**).

#### Generation of Promacrobody Nb fusions (PMB)

A synthetic gene encoding the maltose binding protein fragment from the promacrobody (PMB) system^29^ was ordered and inserted into each pET22b vector used for expression of the above nanobodies using Gibson assembly. Primers are listed in **Table S1**. PMBs were purified using the same method as nanobodies.

### Nanobody Functionalization

#### Sortase mediated ligation

Sortase mediated ligation followed a previously published protocol.^25–26^ Briefly, 200 μL of Ni-NTA resin was equilibrated with sortase reaction buffer (50 mM HEPES pH 7.5, 150 mM NaCl, 10 mM CaCl_2_.). Purified sortase A (P94S/D160N/D165A/K196T) (40 μg) bearing a 6xHis tag was added to the equilibrated resin, volume was adjusted to 600 μL, and the mixture incubated on ice for 10 minutes. The mixture was transferred to a 2 mL plastic column, resin was allowed to settle for 5 minutes, and excess liquid was removed from the resin bed by gravity flow. Individual nanobodies bearing the C-terminal sortag were mixed with a GGGYK-SulfoCy5-peptide (21^st^ Century Biochemicals) to a final concentration of 5 μM Nb and 7 μM Cy5-peptide and adjusted to a final volume of 3 mL in sortase reaction buffer. The Nb-peptide mixture was added to the column, and flowthrough collected in 0.5 mL fractions. The column was then washed with 3 mL of 7 μM Cy5-peptide in sortase reaction buffer and flowthrough collected in 0.5 mL fractions. Finally, the column was washed with 2 mL sortase reaction buffer, and flowthrough collected in 0.5 mL fractions. Unreacted protein was eluted with 3 mL elution buffer (50 mM HEPES pH 7.5, 150 mM NaCl, 400 mM imidazole, 10% glycerol). Fractions were analyzed by SDS-PAGE, with images taken in the Cy5 channel prior to staining the gel with colloidal Coomassie dye. Fractions containing labeled Nb (Cy5 Nbs) were pooled and dialyzed against HEPES buffered saline (HBS) (20 mM HEPES pH 7.5, 150 mM NaCl) + 10% glycerol using Slide-A-Lyzer dialysis cassettes with a 10 kDa MW cutoff (Thermo Fisher Scientific).

### Nanobody Complex Characterization

#### SMA-PAGE

membrane proteins in SMALP were diluted to 1 μM in 50 mM HEPES pH 8.0, 150 mM NaCl. Cy5 Nbs was then added at either a 1.2-fold or 0.6-fold molar ratio, and volume was adjusted to 15 μL. Native loading dye consisted of 1 mg/mL bromophenol blue, 20 mM Tris pH 8.0, and 50% (v/v) glycerol. Native gels were run using pre-cast 4-20% gradient gels (Bio-rad) using cold 25 mM Tris pH 8.8, 192 mM glycine running buffer. Gels were run at 200 V for 34 minutes at room temperature. Prior to Coomassie staining, gels were imaged on a GelDock at the Cy5 channel (BioRad). Gels were subsequently stained and imaged again.

#### Biolayer Interferometry (BLI)

All BLI experiments were performed using PBSA supplemented with 0.05% Tween-20 (BLI Binding buffer). Data were collected on an Octet RED96 instrument (Sartorius). After NiNTA regeneration as detailed in the manual,^33^ NiNTA-functionalized tips were hydrated for 10 minutes in BLI binding buffer, and a 60 second baseline was measured. Tips were loaded with Nanobodies bearing a C-terminal 6xHis tag diluted to 100 nM in BLI binding buffer for 120 seconds. After loading, a second baseline measurement was collected in BLI binding buffer for 60 seconds. Afterwards, tips were dipped into solutions of bait protein in SMALP at various concentrations (1500 nM to 125 nM) to measure association for 600 seconds. Tips were transferred back to BLI binding buffer for 600 seconds to measure dissociation. Control measurements at each concentration of bait protein in SMALP were recorded to account for baseline drift during association and dissociation. All data were collected in biological duplicates. Baselines were subtracted, and association and dissociation curves were fit using GraphPad Prism (GraphPad Software, San Diego CA). A 1:1 binding mode was fit using the “associate then dissociate” model for PglI, Wzx, and PglB Nbs. Due to the high affinity of the WbaP Nb, the 1-phase association and dissociation curves were fit separately with GraphPad Prism and the K_D_ was calculated manually.

#### Size Exclusion Chromatography (SEC)

##### S. enterica WbaP

For measurement of Nb binding, 150 μL of purified *S. enterica* WbaP in SMA200 at 15.3 μM was mixed with purified WbaP Nb to a final molar ratio of 1:1.2 WbaP:Nb. The complex was incubated on ice for 30 minutes and injected onto an Enrich SEC 650 10/300 (Bio-RAD) column on a AKTA system pre-equilibrated with 25 mM HEPES pH 8.0, 150 mM NaCl. Peak fractions were analyzed by SDS-PAGE to confirm co-elution of Nb and WbaP.

For measurement of PMB binding 40 µM of purified WbaP was incubated with 48 µM PMB at a ratio of 1:1.2 WbaP:PMB fusion at a total volume of 500 µL. The complex was incubated on ice for 60 minutes and injected onto a Superose 6 10/300 GL (Cytiva) column pre-equilibrated with 25 mM HEPES pH 8.0, 150 mM NaCl. Peak fractions were analyzed by SDS-PAGE to confirm co-elution of WbaP and PMB.

##### C. concisus PglI

A 17 µM sample of purified *C. concisus* PglI in AASTY 6-45 was incubated with 20.4 µM PglI Nb and 0.27 µM Cy5-labeled PglI Nb at a total volume of 100 µL. The Cy5-labeled PglI Nb was used as an additional marker for tracking during SEC. The complex was incubated on ice for 30 minutes and injected onto a Superose 6 Increase 3.2/300 (Cytiva) column on an AKTA micro system pre-equilibrated with 25 mM HEPES pH 8.0, 150 mM NaCl. Peak fractions were analyzed by SDS-PAGE to confirm co-elution of Nb and PglI.

##### N. gonorrhoeae PglB

A 40 µM sample of purified *N. gonorrhoeae* PglB in AASTY 6-50 was incubated with 48 µM PMB at a ratio of 1:1.2 PglB:PMB at a total volume of 500 µL. The complex was incubated on ice for 60 minutes and injected onto a Superose 6 10/300 GL (Cytiva) column on an AKTA Pure system pre-equilibrated with 25 mM HEPES pH 8.0, 150 mM NaCl. Peak fractions were analyzed by SDS-PAGE to confirm co-elution of PglB and PMB.

## Supplementary Material Description

Supplementary information includes a table of the primers used for cloning the Nbs (Table S1), SDS-PAGE showing purification of proteins (Figure S1), binding data to show specificity of the yeast population (Figure S2), FACS sorting of WbaP population (Figure S3), titration of binding of WbaP population (Figure S4), analysis of selected clones binding (Figure S5), Nb sequence alignment (Figure S6), SDS-PAGE showing purification of Nbs (Figure S7), nanoDSF data for Nbs (Figure S8), graphic of sortase mediated ligation (Figure S9), SDS-PAGE of Cy5 conjugated Nbs (Figure S10), and non-denaturing SMA-PAGE of WbaP and Wzx (Figure S11).

## Supporting information

Supplement containing Figs S1-S11, Table S1

## Acknowledgments

This work was supported by NIH R01 GM039334 and R01 GM131627 (to KNA and BI), F32 GM149160 (to HK) F32 GM134576 (to GJD). The Octet RED96 instrument is housed and supported by the MIT Biology Biophysical Instrumentation facility. Selected figures were created with biorender.com.

