## Supplement containing Figs S1-S11, Table S1 for "Selection of nanobodies against liponanoparticle-embedded membrane proteins by yeast surface display"

**Table S1:** Primers used for Nbs.

| Primer name | Primer sequence 5’ -> 3’ |
|---|---|
| **Pet22_Nb_FWD** | CCAGCCGGCGATGGCCCAGGTGCAGCTGCAGG |
| **Pet22_Nb_REV** | GTGGTGGTGGTGCTCGAGGCTGCTCACGGTCAC |
| **Pet22_PMB_fwd** | CACGGTCACCTGGGTGC |
| **Pet22_PMB_rev** | CTCGAGCACCACCACCACCA |
| **Nb_Sortase_primer_1** | ACTGCCGGAAACCGGCGGCCTCGAGCACCACCACCAC |
| **Nb_Sortase_Primer_2** | CCCCCACTCCCGCTACCACCGCTGCTCACGGTCACCTG |

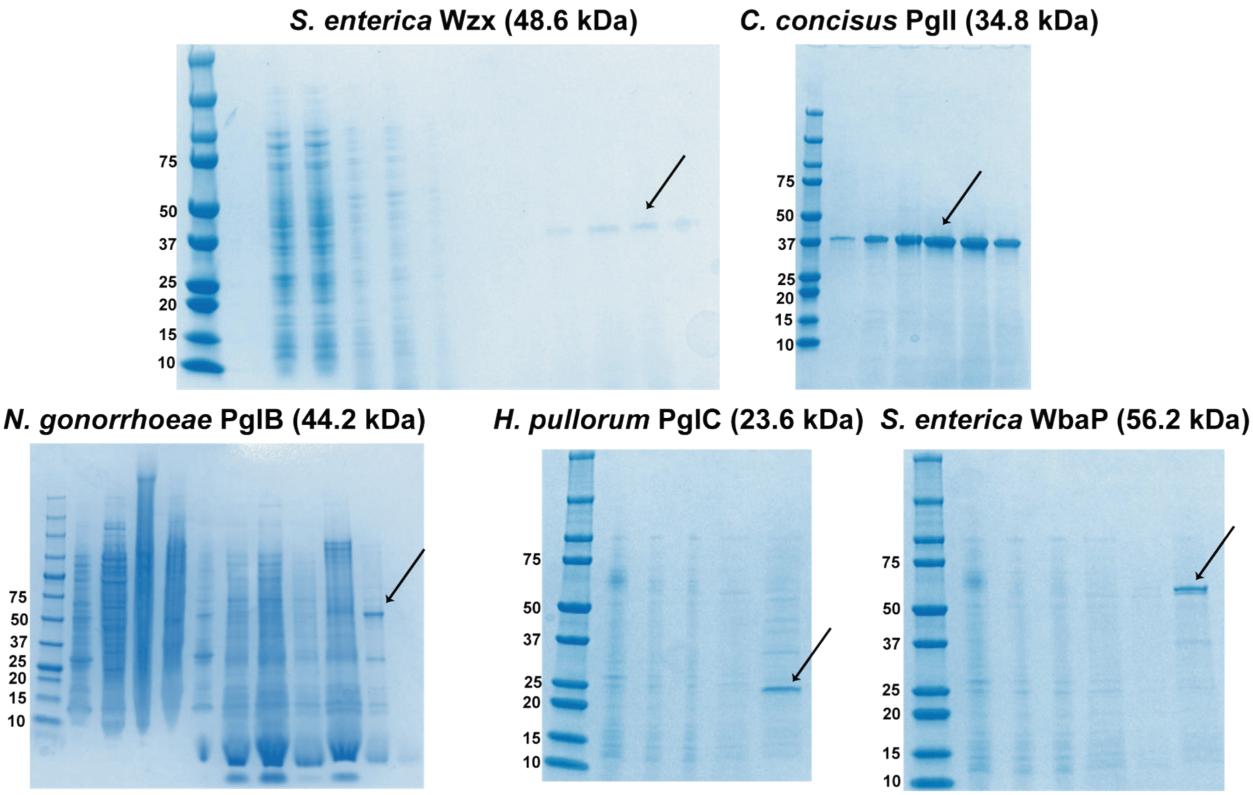


**Figure S1**. SDS-PAGE showing prurification of enzymes in various polymers with their predicted molecular weight listed. Black arrow points at the band of interest.

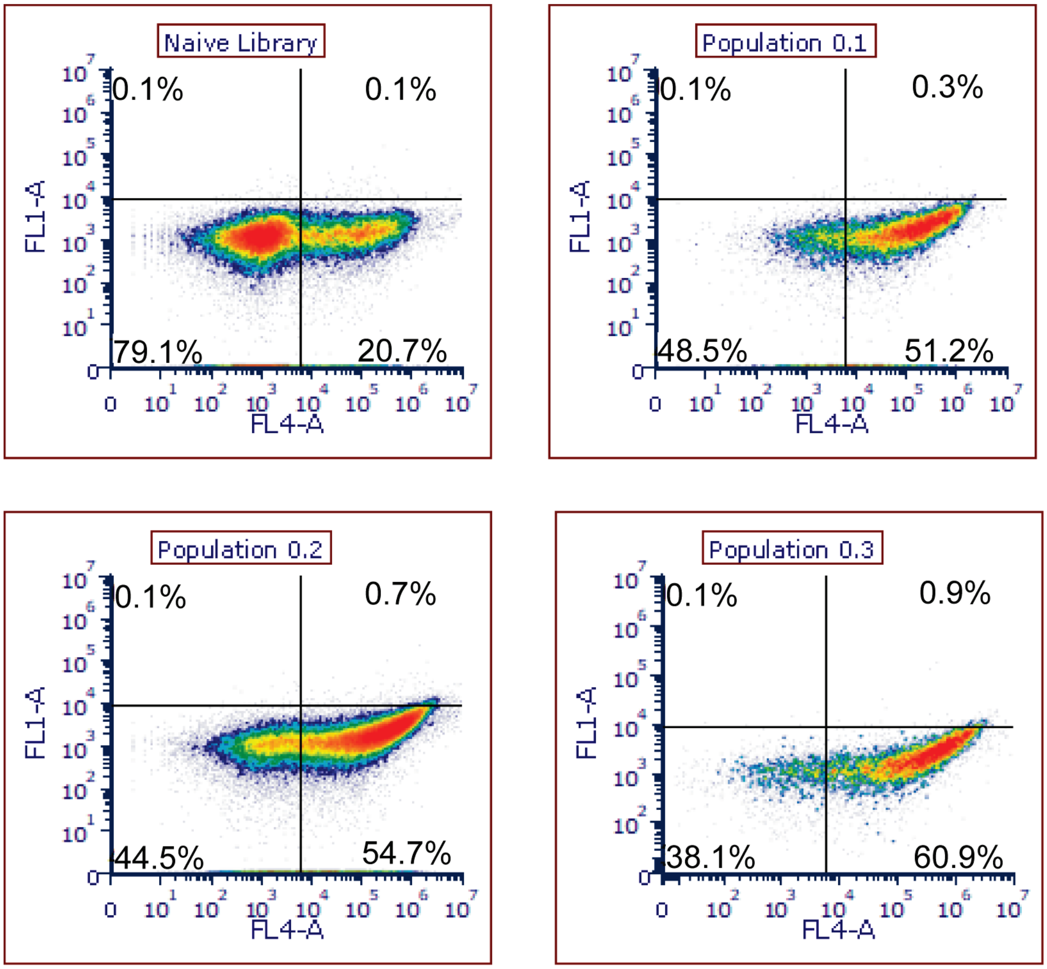


**Figure S2**. Assessment of selected WbaP yeast population specificity. Each selected yeast population from the naïve library to population 0.3 was assessed by analytical flow cytometry for binding to *A. hydrophila* WecP, a WbaP homolog with 36% identity. Minimal enrichment was observed, indicating these populations were specific for WbaP, and were unlikely to bind the shared dual-strep tag or SMALP components.

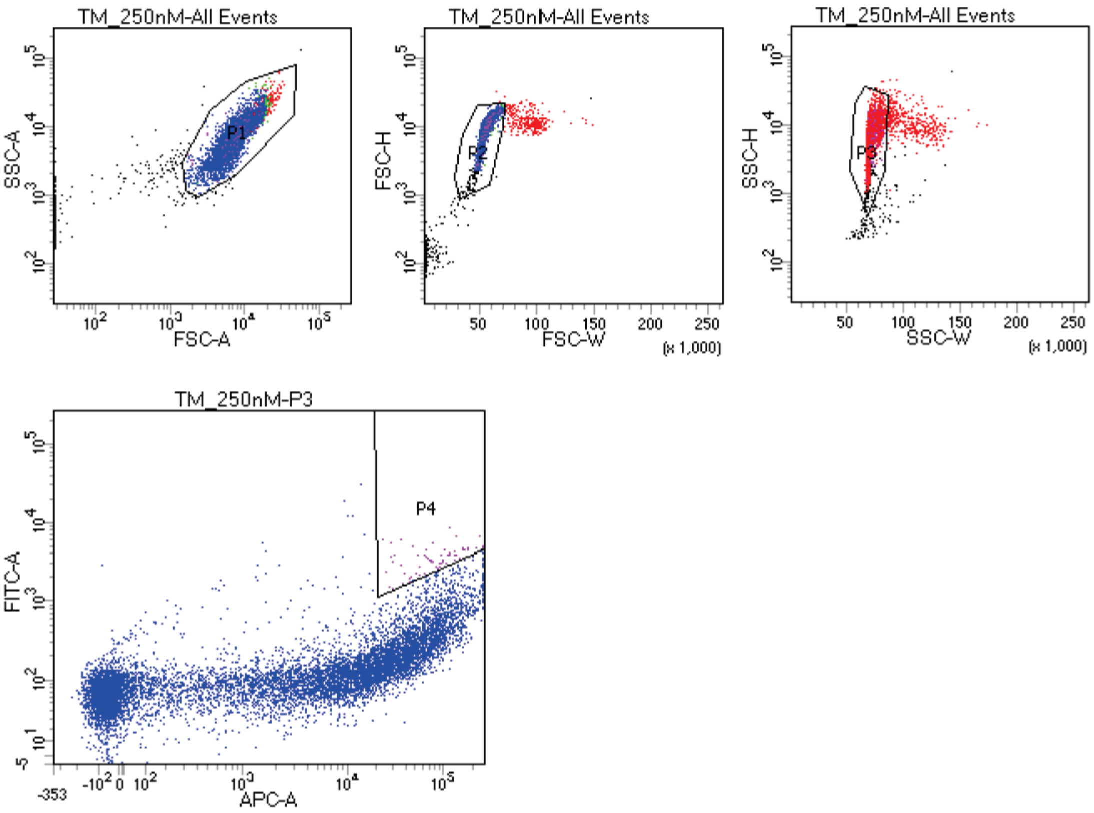


Tube: TM_250nM

| Population | #Events | %Parent | %Total |
|---|---|---|---|
| All Events | 10,632 | #### | 100.0 |
| P1 | 10,371 | 97.5 | 97.5 |
| P2 | 10,048 | 96.9 | 94.5 |
| P3 | 9,959 | 99.1 | 93.7 |
| P4 | 66 | 0.7 | 0.6 |

**Figure S3.** Fluorescence activated cell sorting of WbaP population 0.3. SSC: side scatter, FSC: forward scatter, FITC-A: fluorescence channel detecting Alexafluor488-tagged streptactinXT, APC-A: fluorescence channel detecting Alexafluor647-tagged nanobodies on yeast surface. 20,377 were sorted using gate P4 to create population 0.4.

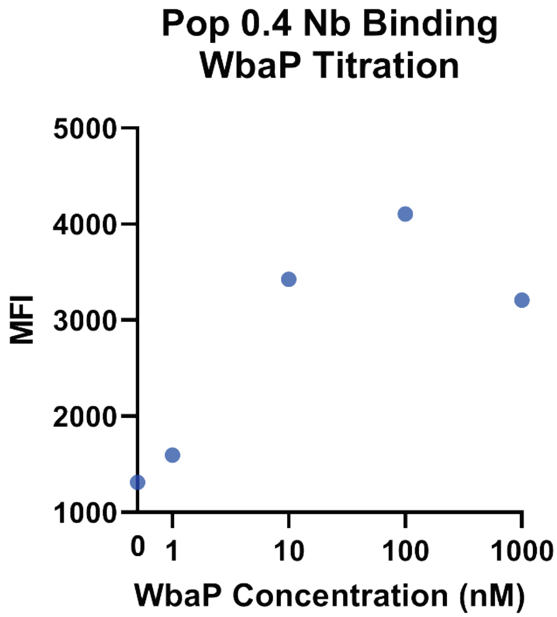


**Figure S4.** Titration of WbaP binding to population 0.4 by analytical flow cytometry. Population 0.4 exhibited maximum binding to 100 nM WbaP, with a decrease in binding at 1μM WbaP.

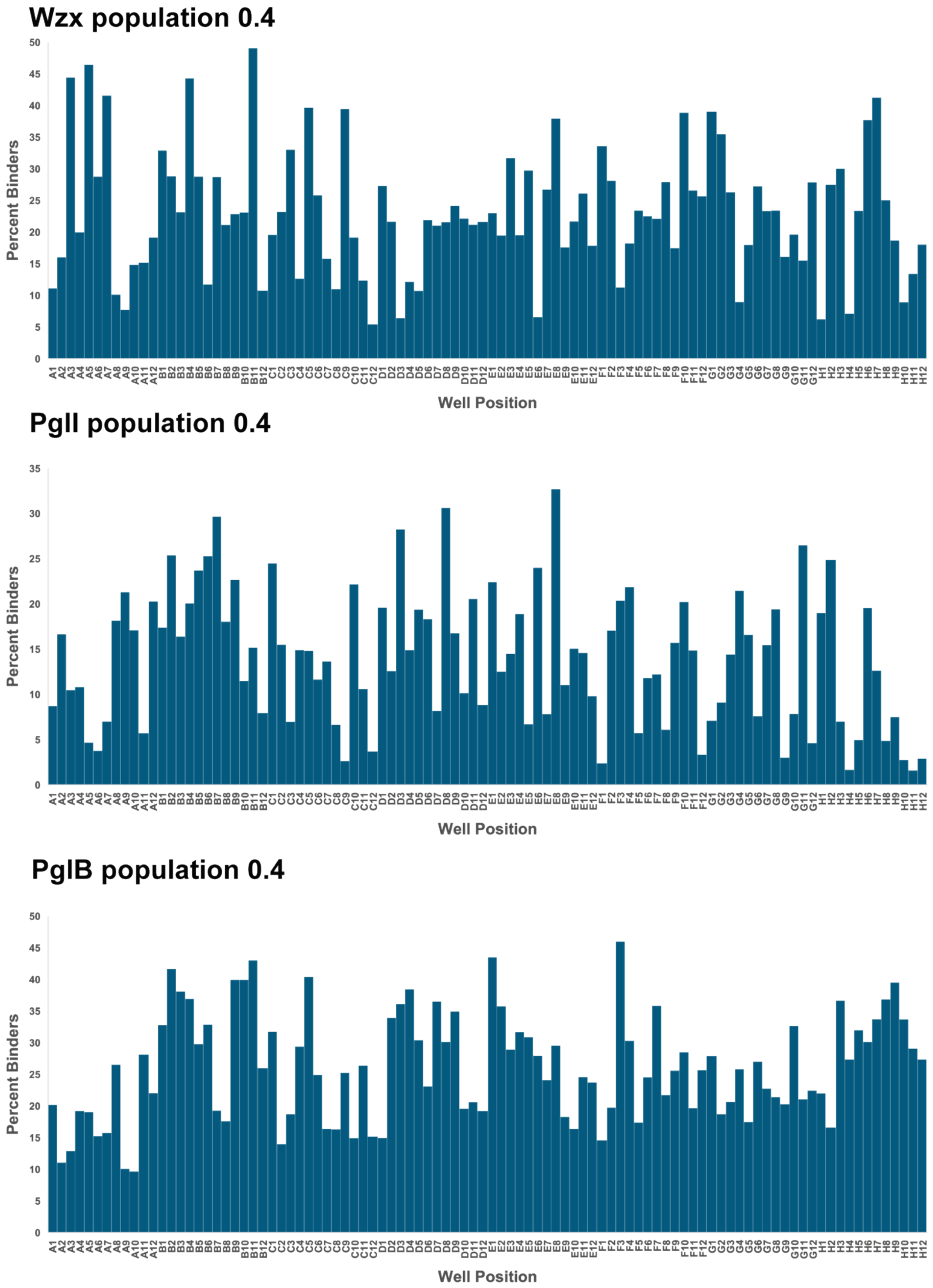


**Figure S5**. Analysis of selected individual clones from population 0.4 of Wzx (top), PglI (middle), and PglB (bottom) based on highest relative percentage binder.

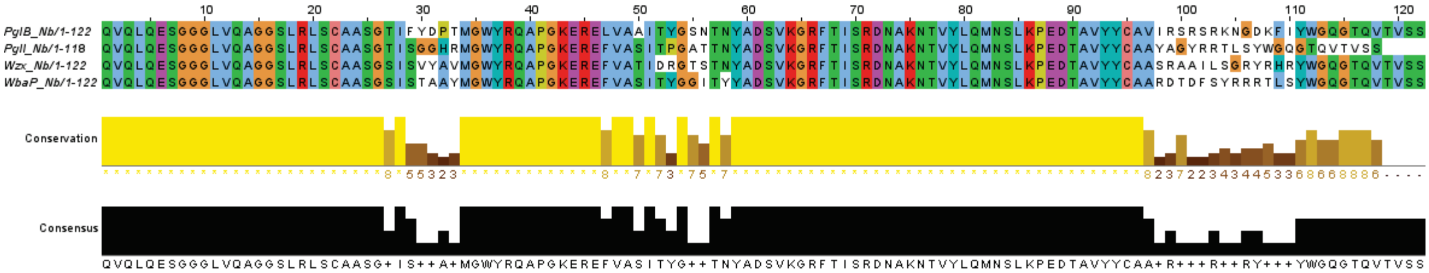


**Figure S6**. Sequence alignment of Nbs.

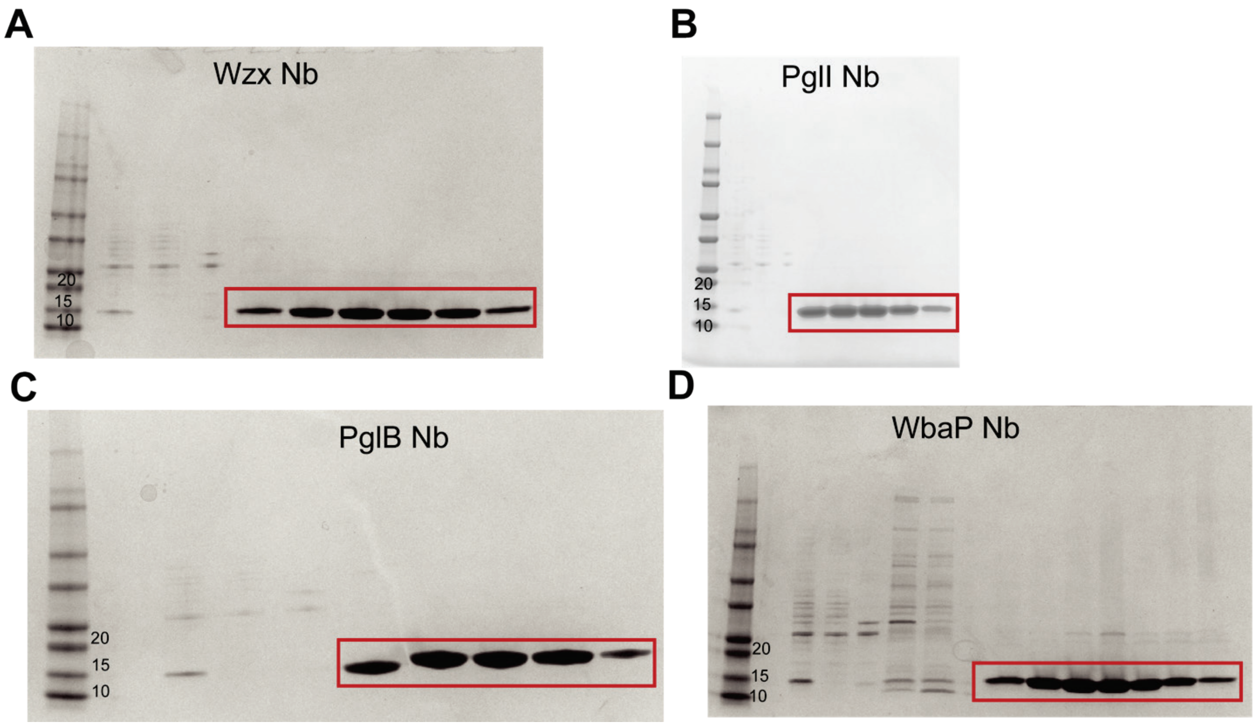


**Figure S7**. SDS-PAGE gels (4 - 20% gradient) showing purification of Nbs, (**A**) Wxz, (**B**) PglI, (**C**) PglB, and (**D**) WbaP. The Nb is boxed in red.

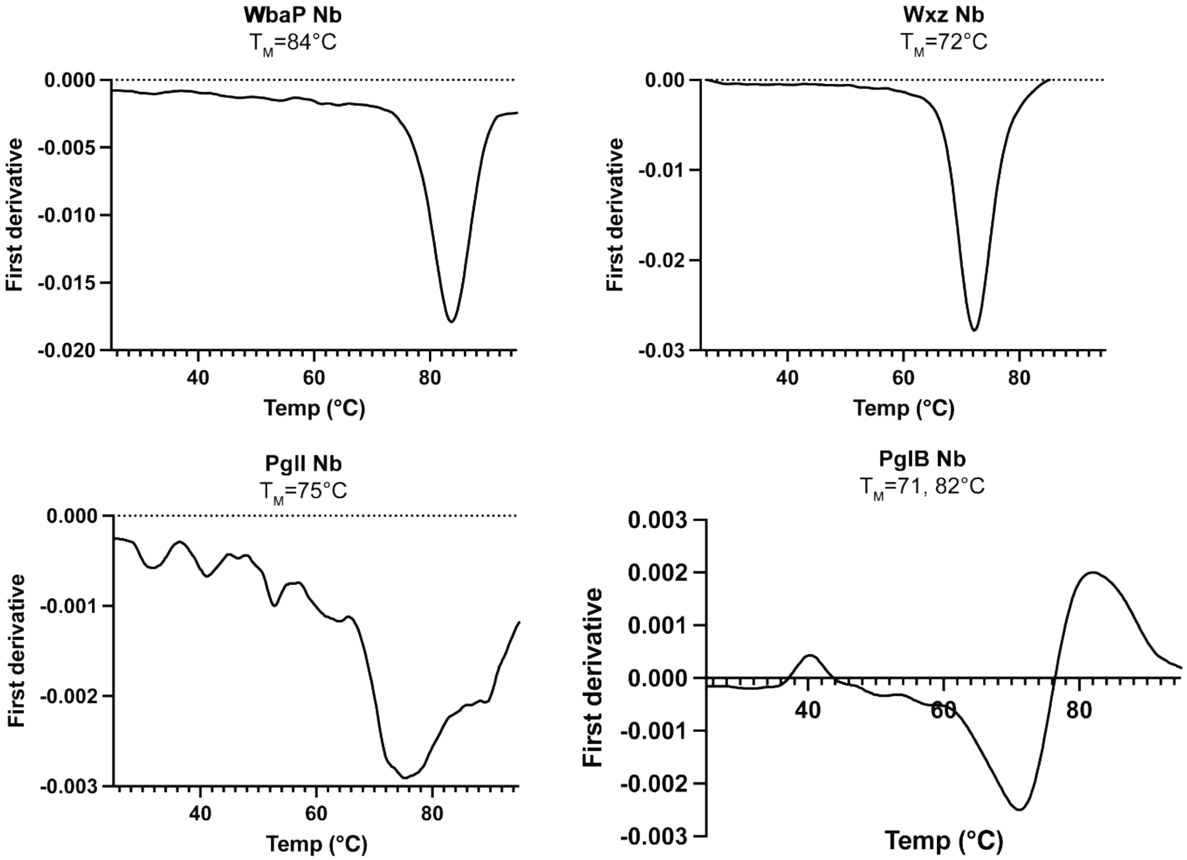


**Figure S8.** NanoDSF of Nbs. The first derivative of the ratio of 350/330 nm is displayed with the associated $T_M$ listed.

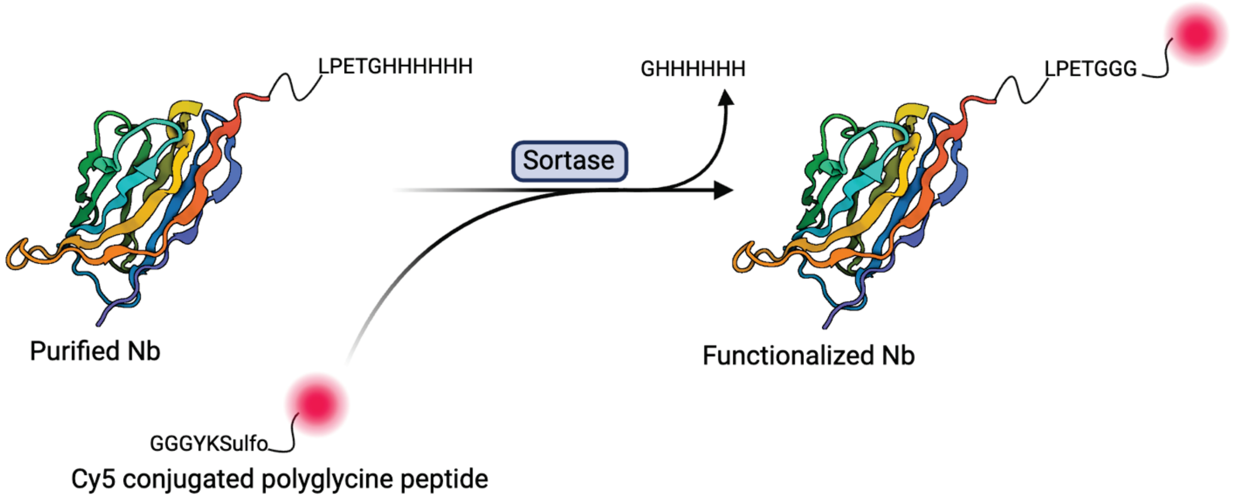


**Figure S9**. Sortase mediated ligation (SML) to generate functionalized Nb probes. Nb model is represented by PDB ID 5VNV.

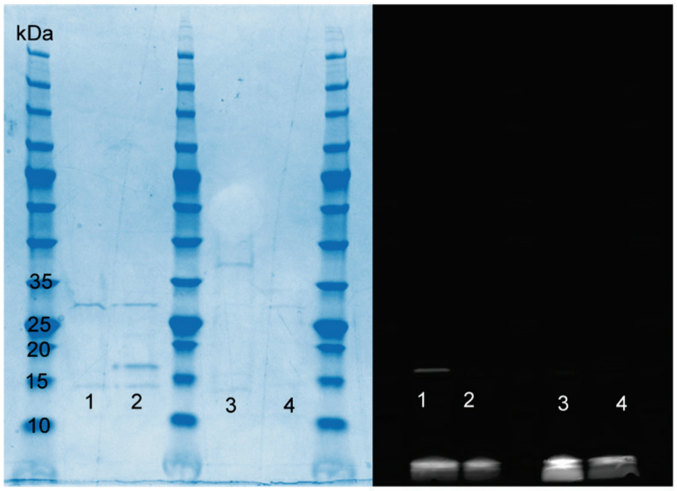


**Figure S10**. Cy5 conjugated Nbs. (left) SDS-Page of Cy5 Nbs. The band at 15 kDa is the monomeric form of the Nb and the higher band is likely the dimer. (right) Cy5 fluorescence of the SDS-Page to the left. The bottom band is excess dye. The lanes are as follows: WbaP Nb (1), PglB Nb (2), PglI (3), and Wzx (4). The band of interest is above the number.

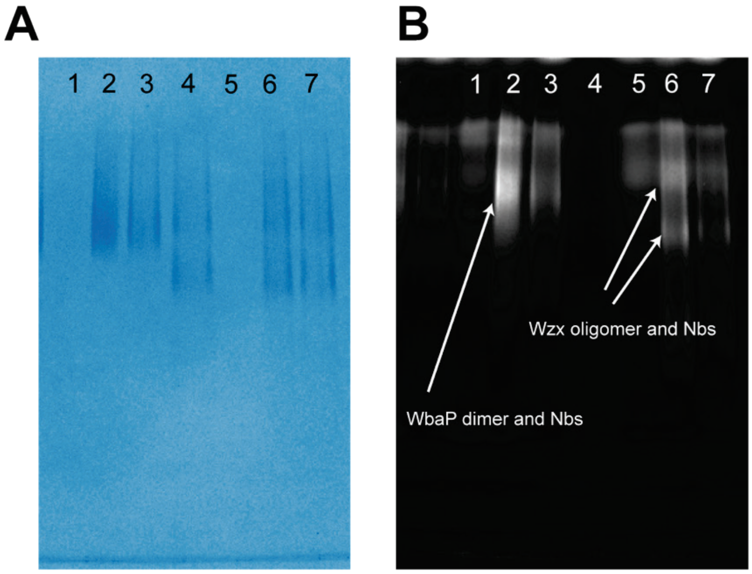


**Figure S11**. SMA-PAGE (non-denaturing) results for WbaP and Wzx. (**A**) Coomassie stained native gel. (**B**) Cy5 imaged native gel. Lanes: WbaP Nb (1), WbaP + 1.2x Nb (2), WbaP + 0.6x Nb (3), Wzx (4), Wzx Nb (5), Wzx + 1.2x Nb (6), and Wzx + 0.6x Nb (7).
